# *In vivo* aberration measurement and correction for ultrafast FACED two-photon fluorescence microscopy of the brain

**DOI:** 10.64898/2026.02.06.704504

**Authors:** Jun Zhu, Ryan G. Natan, Jian Zhong, Iksung Kang, Na Ji

## Abstract

Ultrafast two-photon fluorescence microscopy (2PFM) based on free-space angular-chirp-enhanced delay (FACED) enables megahertz line scanning and kilohertz frame rates for *in vivo* brain imaging. However, optical aberrations from the imaging system and brain tissue degrade spatial resolution, signal, and contrast at depth. Here we integrate adaptive optics (AO) with FACED 2PFM to achieve synapse-resolving ultrafast imaging in the living mouse brain. Because FACED generates a one-dimensional array of temporally delayed, spatially separated excitation foci at 1 gigahertz, we developed a focus-averaging, frequency-multiplexed aberration measurement method that simultaneously measures and corrects the average aberration across all FACED foci using a segmented deformable mirror. We validated the accuracy of our method in correcting both system and artificial aberrations. When applied to *in vivo* morphological imaging of the mouse brain, AO enhances resolution, signal, contrast of dendritic shafts, spines, and boutons. Functionally, AO improves cerebral blood flow imaging by increasing plasma signal and kymograph contrast over large fields of view; when used for glutamate imaging, it amplifies transient amplitudes and reveals visually evoked glutamate release that were undetectable without correction. Together, these results establish AO-FACED 2PFM as a powerful approach that combines ultrafast imaging with high spatial resolution for structural and functional imaging in the living mouse brain.

## Introduction

Understanding brain function requires measuring neuronal structural and functional dynamics *in vivo*, ideally at millisecond temporal and subcellular spatial resolution to capture fast physiological events at individual synapses. With submicron lateral resolution, two-photon fluorescence microscopy (2PFM)^1,2^ is commonly used for *in vivo* brain imaging, but its imaging speed over a field of view (FOV) of hundreds of microns is limited by the mechanical inertia of galvanometer and resonant mirrors to tens of frames per second (fps)^3,4^.

To overcome these speed limitations, we previously developed an ultrafast 2PFM technique leveraging free-space angular-chirp-enhanced delay (FACED)^5–7^. Utilizing a cylindrical lens and a pair of highly reflective, quasi-parallel mirrors to transform an input laser pulse into a one-dimensional (1D) array of spatially separated and temporally delayed excitation foci at the objective focal plane, a FACED module enables line scanning at the laser repetition rate. We have demonstrated FACED 2PFM with line-scan rates up to 4 MHz and frame rates up to 3000 fps^5^, which has enabled ultrafast imaging of cerebral blood flow^6,7^ as well as large-scale voltage and calcium imaging in the mouse brain^7^, while maintaining the spatial resolution and imaging depth characteristics of conventional 2PFM.

Despite these advancements in imaging speed and throughput, a persistent challenge in *in vivo* brain imaging is optical aberration^8–10^. As two-photon excitation light propagates through optically heterogeneous brain tissues, its wavefront becomes increasingly distorted with imaging depth. These sample-induced aberrations enlarge the excitation focal volume and diminish focal intensity, leading to the degradation of image resolution, signal, and contrast. In the mouse brain, optical aberrations, if uncorrected, limit the ability to visualize and resolve fine features such as synapses.

Adaptive optics (AO) measure and correct optical aberrations, thereby offering a powerful solution for maintaining resolution deep within the brain *in vivo*^8–10^. Once the wavefront distortion experienced by the excitation light is known, a corrective wavefront can be applied via a phase correction device such as a deformable mirror to the excitation light, which cancels out the sample-induced aberrations and recovers a diffraction-limited excitation focus. Various AO implementations exist, with indirect wavefront sensing methods particularly well suited for opaque biological media^9,11,12^.

Here, to achieve both high temporal and high spatial resolution for *in vivo* brain imaging, we integrated AO with an ultrafast FACED 2PFM system. The unique implementation of FACED, which generates a 1D array of excitation foci at 1 GHz point-scanning rate, necessitated a specialized AO approach capable of measuring and correcting aberrations across all FACED foci simultaneously. Accordingly, we developed a focus-averaging frequency-multiplexed aberration measurement and correction method. We validated this method by correcting system and artificially introduced aberrations. We then applied it to achieve high-resolution structural visualization of dendrites and dendritic spines, ultrafast blood flow measurements, and glutamate imaging of synaptic structures in the living mouse brain.

## Results

### Adaptive-optical FACED (AO-FACED) 2PFM with focus-averaging multiplexed aberration measurement

We built a two-photon fluorescence microscope with a FACED module for ultrafast scanning and an AO module for aberration measurement and correction (**Fig. 1a**). The FACED module^13^ was composed of a cylindrical lens (CL), a polarizing beam splitter (PBS), a quarter-wave plate (QWP), and a pair of highly reflective mirrors (FM1, FM2) with a relative angle of α (**Fig. 1a**). Through retro-reflections, the FACED module converted each pulse of a femtosecond laser into a set of time-delayed pulses propagating at slightly different angles as follows.

**Fig. 1.**
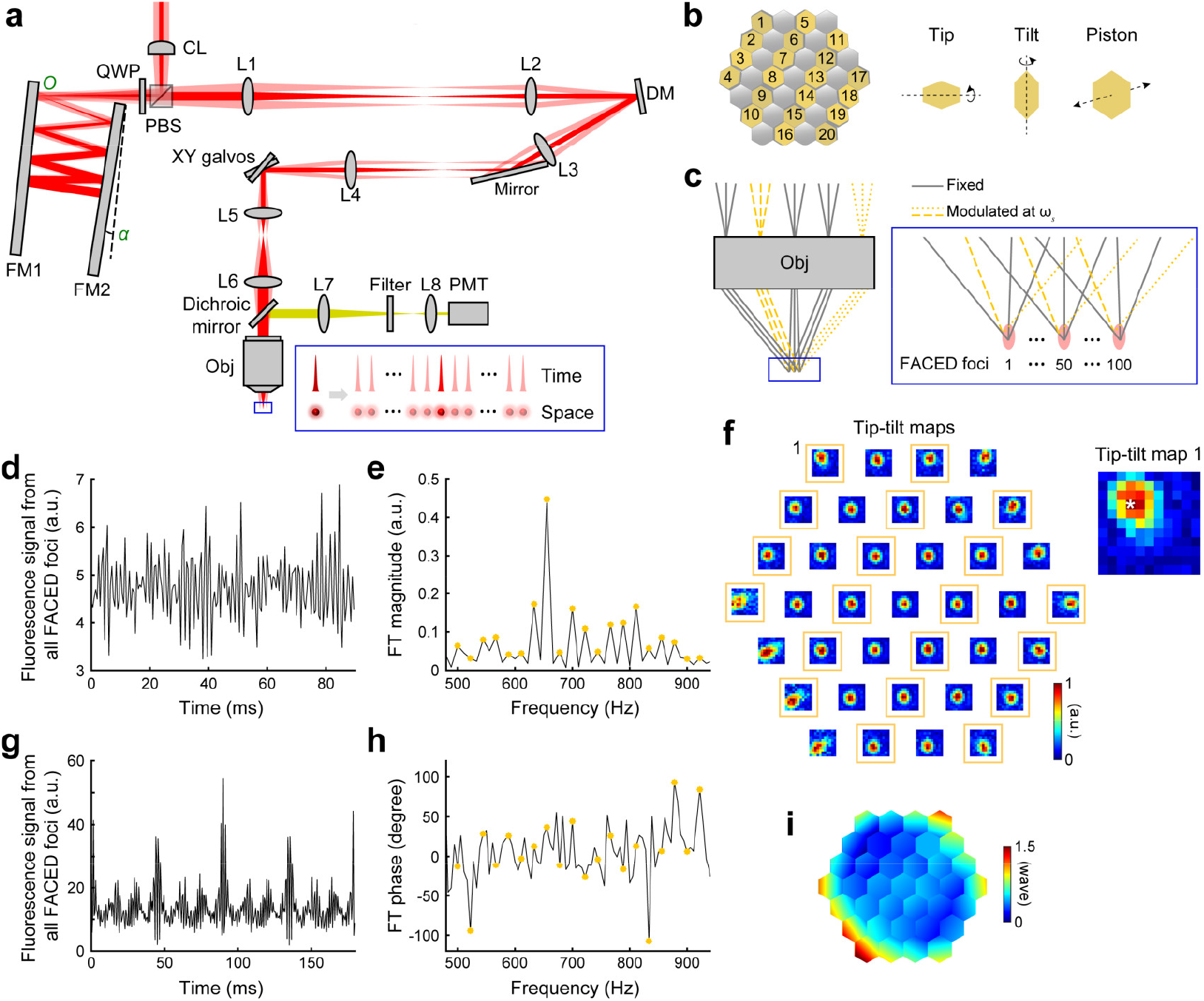
An AO-FACED two-photon fluorescence microscope with focus-averaging frequency-multiplexed aberration measurement. (**a**) Schematic of the AO-FACED system. CL: cylindrical lens; PBS: polarizing beam splitter; QWP: quarter-wave plate; FM: FACED mirror; L: lens; DM: deformable mirror; Obj: objective; PMT: photomultiplier tube. Inset: schematic of the FACED focal array in time and space. (**b**) A 37-segment DM with tip, tilt, and piston control. (**c**) FACED output enters the objective at different angles, forming an array of foci (pink ellipses, inset). During aberration measurement, the phases of some FACED beamlets are modulated at distinct ω_s_’s (gold dashed lines; gold segments in **b**) and the rest are fixed (gray solid lines; gray segments in **b**). (**d**) Two-photon fluorescence signals integrated from all FACED foci during modulation of DM segments at a specific set of tip/tilt angles. (**e**) Fourier Transform (FT) of **d** with FT magnitudes at ω_s_’s indicated by gold dots. (**f**) 2D interference maps of modulated segments (gold boxes) are generated from FT magnitudes at all tip/tilt angle combinations. Switching the modulated and fixed segments and repeating steps in **d**,**e**, interference maps of all segments are obtained. The tip/tilt correction for each segment is determined to be the angles providing maximal FT magnitude/interference (white asterisk in zoomed-in view of Segment 1 map). (**g**) Two-photon fluorescence signals integrated from all FACED foci during phase modulation of half of DM segments. (**h**) FT of **g**, with FT phases at ω_s_’s indicated by gold dots and corresponding to the piston offsets required for each modulated segment. (**i**) Corrective wavefront on DM.

The cylindrical lens focused an excitation laser pulse in 1D, which reflected off the PBS and arrived at the mirror pair. The focus of the cylindrical lens was located at the first mirror that the excitation pulse encountered (entrance *O* on FM1; **Fig. 1a**), after which the pulse reflected between the two mirrors, with the incidence angle reduced by α at each reflection. When the incidence angle for a portion of the pulse became zero, that portion of the pulse was retro-reflected (e.g., dark-red path, **Fig. 1a**). Because of 1D focusing by the cylindrical lens, different portions of the laser pulse had different initial incidence angles and became retro-reflected after distinct number of reflections. As a result, a single input pulse became a series of time-delayed output pulses with distinct propagation directions. The quarter-wave plate, traversed twice by light, rotated the polarization by 90 degrees, enabling the output pulses to transmit through the polarizing beam splitter and enter the microscope. A series of lenses (L1-L6) conjugated *O* to the midplane between a pair of galvanometer mirrors (XY galvos) and the back focal plane of a microscope objective (Obj). These pulses then formed a 1D array of excitation foci that were spatially separated and temporally delayed at the objective focal plane, that is, a line scan (zoomed-in schematic, blue box, **Fig. 1a**). Without utilizing active scanning, the line-scan rate of this all-optical and passive module equals the repetition rate of the laser, which can exceed MHz^5–7^.

With a 1035-nm femtosecond laser operated at 1-MHz repetition rate as the input and the FACED mirror pair placed 150 mm apart at α of 0.0125°, the FACED module generated a 1D array of 100 excitation foci with 1 ns delay and 0.8 μm spacing along the X axis at the focal plane of a 25× 1.05-NA water-dipping objective, achieving line scanning at 1 MHz. The Y galvo then scanned the FACED foci perpendicularly along the Y axis to acquire a FACED FOV. For experiments described here, we scanned a FACED FOV of 80 μm × 400 μm or 80 μm × 200 μm at 1.54 kHz, or 80 μm × 180 μm at 0.94 kHz. When needed, the X galvo tiled the FACED FOV laterally to further enlarge the scanning area. Excited two-photon fluorescence was collected through the same objective, collimated and focused by a pair of lenses (L7, L8), spectrally filtered, and then detected by a photomultiplier tube, whose signal was sampled at 10 gigasamples per second using a high-speed digitizer (**Fig. S1 and Methods**).

The AO module consisted of a 37-segment deformable mirror with tip, tilt, and piston control (DM, **Fig. 1a,b**) that was conjugated to *O* by L1 and L2. The DM surface was then optically conjugated to the midplane between the XY galvos via L3 and L4 and subsequently to the back focal plane of the objective through a pair of telecentric scan and tube lenses L5 and L6. The output pulses from the FACED module reflected off the DM and entered the back focal plane of the microscope objective at different angles. With the DM surface imaged to the objective back focal plane, adjusting the tip, tilt, and piston values of the DM segments controlled the wavefront of all FACED output pulses simultaneously.

The AO module measured the wavefront distortions experienced by the FACED foci and then canceled these distortions by applying a corrective wavefront of opposite phase to the DM. Because the output pulses from the FACED module traveled along slightly different beam paths through the microscope and within the sample (**Fig. 1a,c**), they experienced different aberrations. However, adjacent FACED foci were temporally separated by 1 ns, whereas the DM can be updated only at ∼10 kHz^14,15^, making it impossible to apply a distinct corrective wavefront for each focus. We therefore corrected the average aberration across the 100 FACED foci by modifying a frequency-multiplexed aberration measurement method previously developed by us^11,12,16,17^ to use the focus-averaged fluorescence signal as input (**Fig. S1** and **Methods**). Because a single aberration correction can improve image quality over hundreds of microns within the mouse brain^12,18^, the aberrations experienced by the FACED foci spanning 80 μm in the sample should be highly similar, making our focus-averaging approach an effective solution.

To obtain diffraction-limited excitation foci, we first determined the tip and tilt angles of 37 DM segments so that the beamlets reflected by these segments maximally overlapped at the focal point. We then determined the piston values of the DM segments so that these beamlets were in phase and constructively interfered at the focal point.

During tip/tilt correction, we fixed half of the DM segments (gray segments, **Fig. 1b**), with the beamlets reflecting off these fixed segments (gray lines, **Fig. 1c**) forming 100 reference FACED foci. We then applied a set of tip and tilt angles to the other half of the DM segments (gold segments, **Fig. 1b**), which changed the location of their corresponding beamlets (gold lines, **Fig. 1c**) at the objective focal plane for all FACED foci (zoomed-in schematic, blue box, **Fig. 1c**). Here, the tip/tilt angles of each segment were chosen randomly from an array of angles that scanned their corresponding beamlets over an 11 × 11 grid with a step size of 1.1 µm, 1.5 µm, or 1.8 µm at the focal plane.

At each set of tip and tilt angles, we modulated the phase of FACED beamlets by varying the piston values of their corresponding DM segments at distinct frequencies ω_*s*_ (*s* = 1, 2, …, 20 for the modulated segments in **Fig. 1b**) while recording the total fluorescence signals generated by all FACED foci (**Fig. 1d**). Fourier-transforming this signal trace, we read out the Fourier transform (FT) magnitude at each modulation frequency ω_*s*_ (gold data points, **Fig. 1e**). If a modulated beamlet did not overlap with its reference focus, varying its phase would not impact the signal strength and the FT magnitude would be zero; if the beamlet maximally overlapped with the reference focus, its FT magnitude would reach its maximal value. In other words, the FT magnitude at ω_*s*_ informed how well the corresponding modulated beamlets overlapped with the reference foci. By repeating this measurement for all 11 × 11 sets of tip/tilt angles, we obtained a 2D tip-tilt map for each modulated segment (gold boxes, **Fig. 1f**).

Switching the fixed and modulated segments, we repeated the above procedure and acquired the tip-tilt maps for all segments (**Fig. 1f**). Applying the tip-tilt angles that gave rise to maximal FT magnitudes to each DM segment (e.g., white asterisk, zoomed-in view, **Fig. 1f**), we ensured that all beamlets maximally overlapped at the objective focal plane.

To determine the piston value of each DM segment so that these beamlets were in phase, we also used frequency multiplexing^17^. Similar to tip/tilt correction, half of the segments were fixed, while the other half were phase-modulated at distinct frequencies ω_*s*_’s. From the integrated fluorescence signal (**Fig. 1g**), the FT phases (gold dots, **Fig. 1h**) at ω_*s*_’s corresponded to the piston offsets applied to the DM segments, so that their corresponding beamlets were in phase and constructively interfered at the focus. Switching the fixed and modulated segments and repeating these procedures, we obtained the final corrective wavefront (**Fig. 1i**) that compensated for optical aberrations and enabled all beamlets to maximally overlap and constructively interfere at the FACED foci. This correction process may be iterated 2-3 times to achieve optimal performance.

### Correction of system and artificial aberrations

We first corrected the aberrations inherent in the microscope due to imperfect optical components and alignment. Parking the FACED foci over 2-µm-diameter fluorescent beads at the center of a 400 μm × 400 μm FOV, we used the focus-averaging frequency-multiplexed aberration measurement method to obtain a corrective wavefront (**Fig. 2a**). For a bead at the FOV center, system aberration correction increased its fluorescence signal by 3.6× and decreased its axial full-width-half-maximum (FWHM) from 3.9 μm to 2.6 μm (**Fig. 2b,c**). Even though the corrective wavefront was measured at the FOV center, we observed an overall improvement in axial resolution throughout this FOV, as indicated by the reduction of the beads’ axial FWHMs (**Fig. 2d**). From this point onwards, system aberration was always corrected when “No AO” images were acquired.

**Fig. 2.**
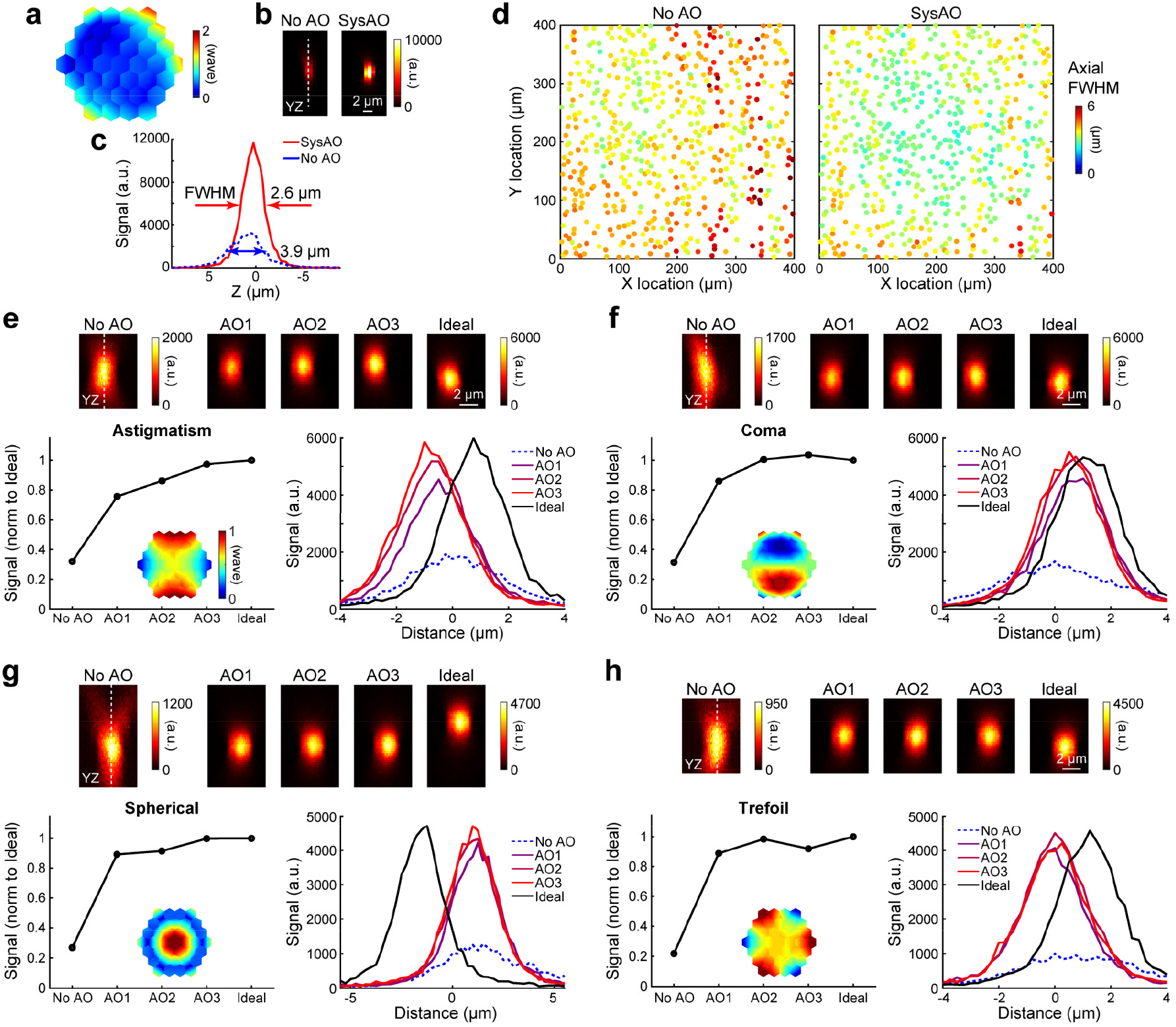
AO-FACED corrects system aberration and artificial aberrations. (**a**) Corrective wavefront for system aberration. (**b**) YZ images of a representative 2-µm-diameter bead without (No AO) and with AO (SysAO) correction. (**c**) Axial signal profiles along dashed white lines in (**b**) labeled with their full-width-at-half-maximum (FWHM) values. (**d**) Axial FWHM values across the FOV without (No AO) and with system aberration correction (SysAO). Pixel size in **b-d**: 0.8 µm × 0.8 µm × 0.25 µm. (**e-h**) Corrections for astigmatism (**e**), coma (**f**), spherical aberration (**g**), or trefoil aberration (**h**) applied to the DM. (Top) YZ images of representative 2-µm-diameter beads acquired without AO (No AO), with iterative AO corrections (AO1, AO2, AO3), and under aberration-free conditions (Ideal); note that different color bars are used for No AO images from other conditions. (Bottom) Maximal signals and axial signal profiles along dashed lines. Pixel size in **e-h**: 0.1 µm × 0.1 µm × 0.25 µm.

To test the accuracy of our method, we applied astigmatism, coma, spherical aberration, or trefoil aberration with a peak-to-valley value of 1 wave to the DM and acquired FACED images of 2-µm beads (“No AO”, **Fig. 2e-h**). Compared with axial images acquired under aberration-free conditions (“Ideal”, **Fig. 2e-h**), the presence of aberration reduced bead signal by 3.1×, 3.2×, 3.7×, and 4.6×, respectively. Beads also became more elongated along the axial direction due to resolution degradation. One round of AO correction substantially improved signal strength and axial resolution (“AO1”, **Fig. 2e-h**); after 3 iterations, image signal and resolution closely resembled the aberration-free condition (“AO3” versus “Ideal”, **Fig. 2e-h**). These results indicate that our approach accurately measures and corrects optical aberrations.

Note that in FACED images, a fluorescence structure can exhibit a “tail” along the FACED/X axis (e.g., bead in **Fig. S2**). This occurred because the temporal delay between adjacent FACED foci was 1 ns, which was shorter than the fluorescence lifetimes of fluorophores in the bead. As a result, fluorophores excited by a FACED focus can emit their fluorescence during data acquisition periods of subsequent FACED foci, leading to elongation of fluorescent features along the FACED axis. We utilized a lifetime deconvolution method^19^ that reassigned photons in these tails to their originating pixels and thereby removed lifetime-induced signal mixing. In this manuscript, we explicitly indicate the use of this deconvolution method in figures where it was applied and provide the raw images in **Supplementary Information**.

### AO improves FACED 2PFM structural imaging in the mouse brain *in vivo*

Next, we used AO to improve *in vivo* structural imaging of neurons in the mouse brain. In a Thy1-GFP line M mouse^20^, we performed focus-averaging frequency-multiplexed aberration measurement to obtain a corrective wavefront (**Fig. 3a**) by placing the FACED foci (spanning the dashed line with arrows, **Fig. 3b**) in line with the cell body of a cortical neuron 280 μm below the dura. Comparing three-dimensional (3D) image stacks acquired without and with the corrective wavefront (maximal intensity projections, **Fig. 3b**), we observed a 1.6× increase in the brightness of the cell body and a general improvement in image quality (**Fig. 3c,d**). Because optical aberrations degrade images of smaller features more severely^11,12^, correcting aberration led to a larger increase in brightness of dendritic spines (e.g., by 2.4× and 2.0× for ROI 1 and 2, **Fig. 3c**) and dendritic shafts (e.g., by 2.0× for ROI 3, **Fig. 3c**). Together with signal increase, their YZ images revealed a substantial improvement in axial resolution (**Fig. 3d**) and a higher contrast for images acquired after aberration correction, with 4.9×, 2.9×, and 4.0× gains in contrast across the line profiles of ROI 1-3 (along dashed white lines, **Fig. 3d** and **Methods**).

**Fig. 3.**
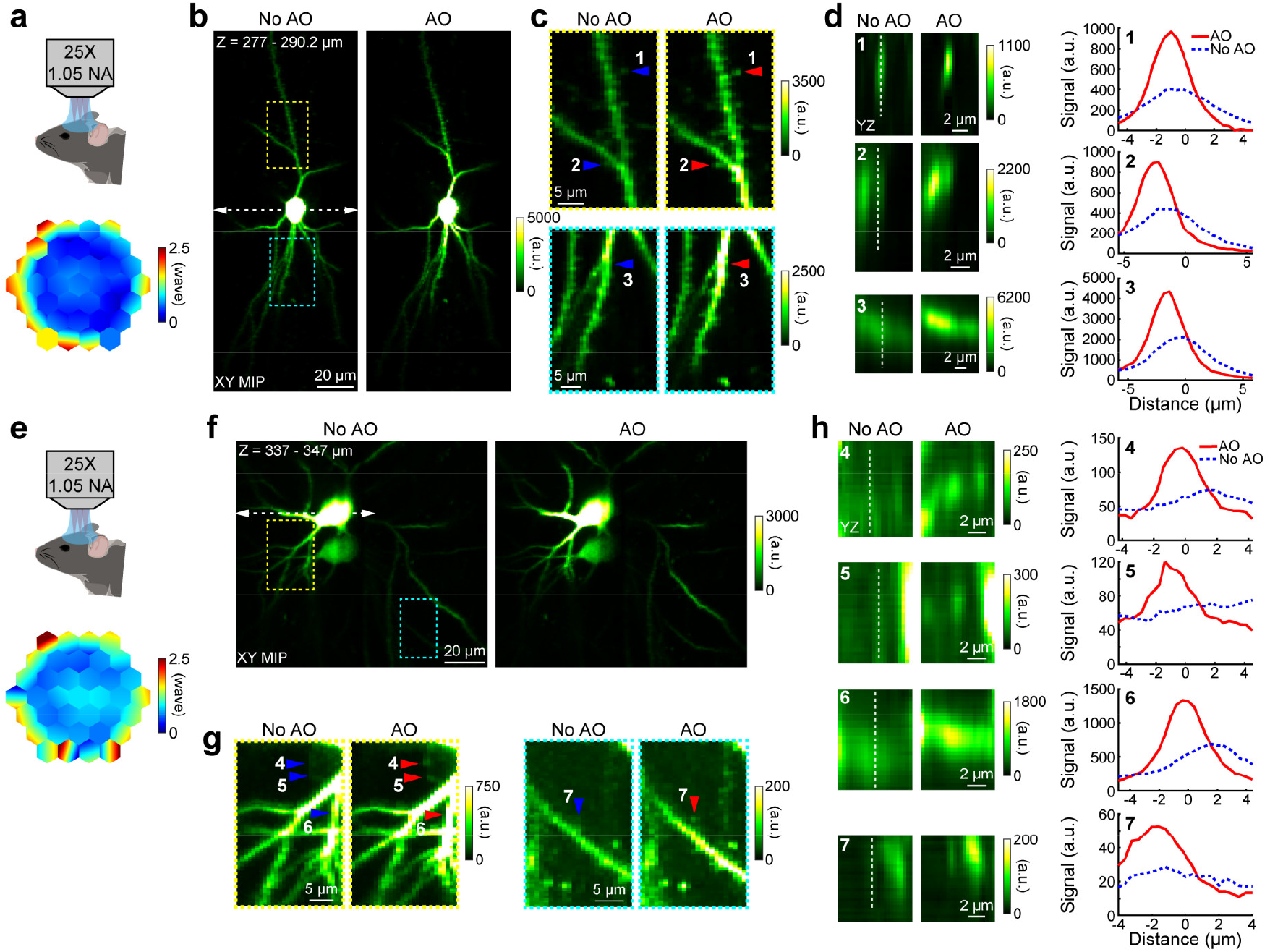
AO improves FACED 2PFM structural imaging in the Thy1-GFP line M mouse brain *in vivo*. (**a**) Corrective wavefront measured from a cell body at 280 μm depth *in vivo*. (**b**) XY maximum intensity projections (MIPs) of 277-290.2 μm depth acquired without and with AO. White dashed line with arrowheads: location of FACED foci during aberration measurement. (**c**) Zoomed-in views of boxed regions in **b**. (**d**) YZ images across dendritic spines (ROI 1,2) and shaft (ROI 3) (indicated by arrowheads in **c**) and their axial signal profiles along dashed white lines. (**e**) Corrective wavefront measured from a cell body at 330 μm depth. (**f**) XY MIPs of 337-347 μm depth acquired without and with AO. (**g**) Zoomed-in views of the boxed regions in **f**. (**h**) YZ images across varicosities (ROI 4,5), dendritic shaft (ROI 6), and spine (ROI 7) (indicated by arrowheads in **g**) and their axial signal profiles along dashed white lines. Pixel size: 0.8 μm × 0.4 µm × 0.4 μm. Excitation power: 151.9 mW. Fluorescence lifetime deconvolution along the FACED/X dimension was applied to XY MIPs (**Fig. S4**).

Aberration correction conducted at a depth of 330 μm over the cell body of another neuron showed similar improvements (**Fig. 3e-h**). AO enabled visualization of fine varicosities (ROI 4,5; **Fig. 3g**), as well as enhanced signal and visibility of dendrites and dendritic spines (e.g., ROI 6,7; **Fig. 3g**). As measured from their YZ images (**Fig. 3h**), AO improved axial resolution and increased their peak signal by 1.6–1.9× and contrast by 2.8–5.1× for the axial profiles of these fine features (along dashed white lines in **Fig. 3h**), enabling them to be clearly resolved in images acquired with AO.

In a wild-type mouse cortex with sparse tdTomato expression, we measured and corrected aberrations using cell bodies 340 μm and 476 μm below dura (**Fig. S3**). Similar to those measured in the Thy1-GFP line M mouse brain, the corrective wavefronts were dominated by spherical aberration (**Fig. S3a,e**). Aberration correction improved visibilities of dendritic shafts (ROI 1,3) as well as synaptic structures such as axonal boutons (ROI 2,6) and dendritic spines (ROI 4,5) (**Fig. S3c,g**). AO substantially decreased the axial sizes of these subcellular features in YZ images and increased their signals by 1.7–2.5× and contrasts by 1.6–3.1× (**Fig. S3d,h**). Together, these results demonstrate that AO-FACED substantially and reliably improved *in vivo* structural imaging deep within the mouse cortex.

### AO improves blood flow imaging in the mouse brain *in vivo*

Having demonstrated the power of AO-FACED for high-resolution structural imaging *in vivo*, we next investigated how aberration correction improved blood flow imaging in the mouse brain. We visualized cortical vessels by labeling the blood plasma via retro-orbital injection of dextran-conjugated Rhodamine B and obtained a corrective wavefront (**Fig. 4a**) using the fluorescence signal of a capillary within the visual cortex (overlaid by a dashed line with arrows, **Fig. 4b**), which increased the time-averaged signal of the capillary by 1.3×. We then measured blood flow by tracking the motion of unlabeled blood cells within the vasculature by acquiring time-lapse images over a 378 μm × 408 μm FOV at 188.7 Hz. We selected ROIs along vessel centerlines (e.g., white dashed lines in **Fig. 4c,d**, the zoomed-in views of two boxed regions in **Fig. 4b**). By stacking the signal profiles extracted from consecutive image frames along each ROI, we generated a kymograph in which individual blood cells appeared as dark stripes, with stripe slopes determined by their flow velocities (e.g., 10-s-long kymographs, **Fig. 4e,f**), from which we determined their mean flow velocity (e.g., using images acquired with AO, 0.55 mm/s for **Fig. 4c,e**; 0.18 mm/s for **Fig. 4d,f**).

**Fig. 4.**
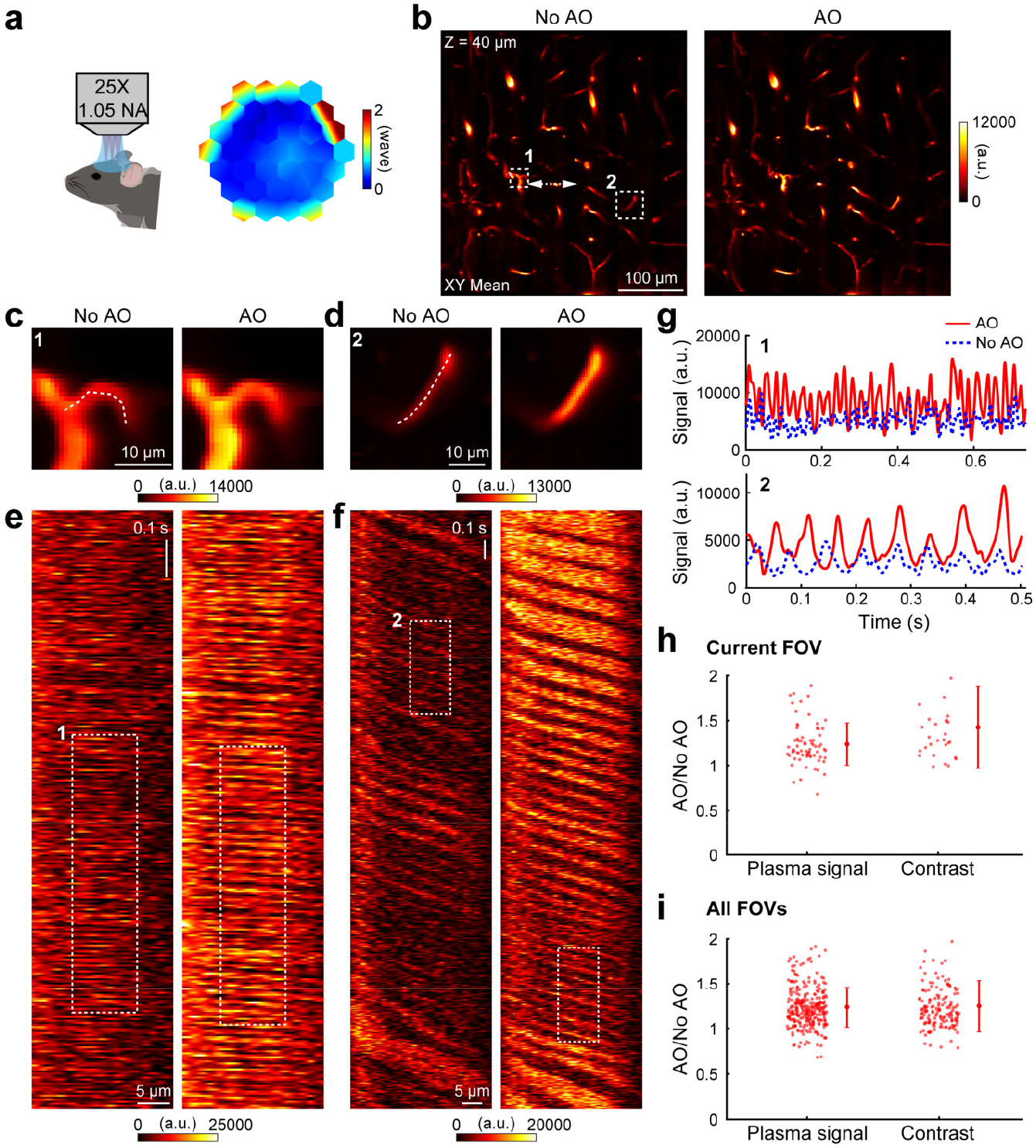
AO improves blood flow imaging in the mouse brain *in vivo*. (**a**) Corrective wavefront measured from a capillary at 40 μm depth in the mouse visual cortex. (**b**) XY images acquired without and with AO. White dashed line with arrowheads: location of FACED foci during aberration measurement. Excitation power: 102.8 mW. Pixel size: 0.8 μm × 0.8 μm. (**c,d**) Zoomed-in views of boxed regions in **b**. (**e,f**) Kymographs along the line ROIs in **b**. (**g**) Average line profiles for the boxed regions in **e** and **f**. (**h**) Scatter plots of AO/No AO ratios for plasma signal (69 segments; left) and line profile contrast (29 segments; right) from capillary segments in **b**. (**i**) Scatter plots for plasma signal (295 segments) and contrast (159 segments) across six FOVs. Error bars: ± s.d.

We selected regions of kymographs with comparable stripe slopes and spacing (e.g., dashed boxes, **Fig. 4e,f**), flattened them, and averaged along the spatial dimension to generate line profiles (**Fig. 4g** and **Methods**). From these line profiles, we quantified the plasma signal, defined as the mean of the highest 20% of signal values in each line profile, and the profile contrast (**Methods**). For both example capillary segments (**Fig. 4g**), AO enhanced the plasma signal by 1.8× and 1.9×, leading to a contrast increase of 1.5× and 1.3×, respectively. Over this FOV, AO significantly increased the plasma signal by 1.24 ± 0.24 (mean ± s.d., 69 segments) and contrast by 1.43 ± 0.45 (29 segments; **Fig. 4h**; p values for one-side tailed t-test: 2.8 × 10^-12^, 1.2 × 10^-5^). Across all six imaged FOVs, the improvement ratios were 1.24 ± 0.22 (295 segments) for plasma signal and 1.25 ± 0.29 (159 segments) for profile contrast (**Fig. 4i**; p values for one-side tailed t-test: 3.2 × 10^-50^, 2.0 × 10^-21^). In summary, AO increases plasma signal and the contrast of kymographs, enhancing blood flow measurement at large depths.

### AO improves FACED 2PFM glutamate imaging in the mouse brain *in vivo*

We also applied AO-FACED to measure glutamate release at synapses of neurons expressing the genetically encoded glutamate indicator iGluSnFR4^21^ in the mouse visual cortex *in vivo*. We corrected cranial window aberration using the signal of a 2-μm-diameter fluorescent bead on the brain surface (**Fig. 5a**). Subsequently, we displayed flash stimulations to the mouse and acquired time-series images of iGluSnFR4-expressing neurons at different depths without and with AO. The MHz-line-scanning rate of FACED enabled us to image a representative 212.8 μm × 181 μm FOV at a depth of 135 μm at 156.3 Hz (**Fig. 5b**). We selected 268 varicosities in the neuronal processes as putative synaptic ROIs within this FOV and quantified how aberration correction impacted their glutamate transients.

**Fig. 5.**
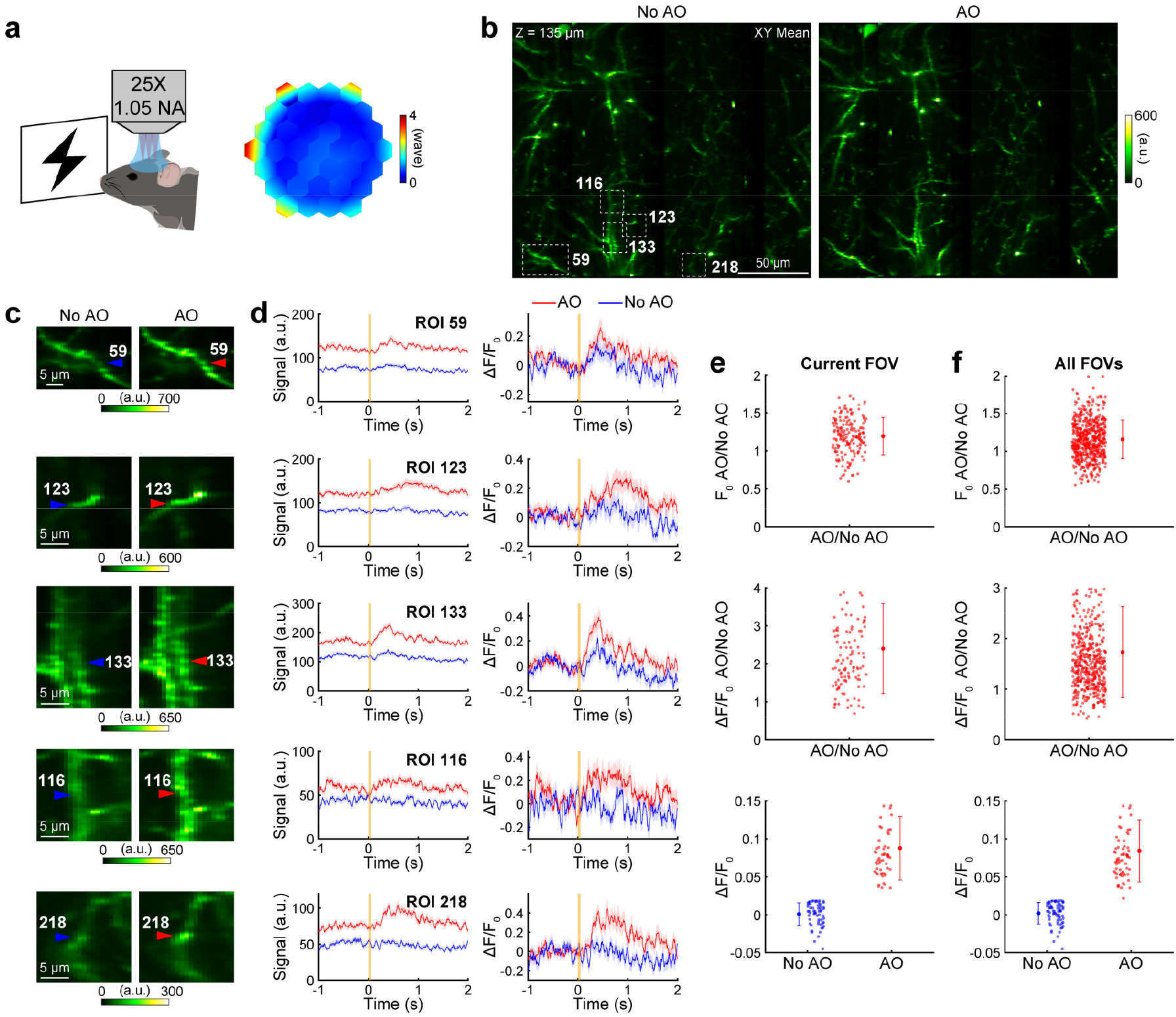
AO improves FACED 2PFM glutamate imaging in the mouse visual cortex *in vivo*. (**a**) Corrective wavefront for cranial window aberration. (**b**) XY images of iGluSnFR4-expressing neurons acquired at 135 μm below dura without and with AO. Pixel size: 0.8 µm × 0.2 µm. Excitation power: 160.5 mW. (**c**) Zoomed-in views of boxed regions in **b**. Arrowheads and numbered ROIs: putative synaptic structures. (**d**) Fluorescence (left) and transient (ΔF/F_0_; right) traces of example ROIs in **c**. 60-trial averages; line and shade: mean and s.d. Orange line: flash stimulation period. (**e**) For FOV in **b**, scatter plots of (top) enhancement ratios of basal fluorescence (F_0_) of 176 visually responsive ROIs, (middle) enhancement ratios of glutamate transient magnitudes (ΔF/F_0_) of the 117 ROIs with mean ΔF/F_0_ ≥ 0.02 under No AO condition, (bottom) mean ΔF/F_0_ measured without and with AO for the 59 ROIs with mean ΔF/F_0_ < 0.02 under No AO condition. (**f**) Scatter plots for 572 visually responsive ROIs, 500 ROIs with mean ΔF/F_0_ ≥ 0.02 under No AO condition, and 72 ROIs with mean ΔF/F_0_ < 0.02 under No AO condition across four FOVs. One-side tailed t-test for whether the improvement ratios significantly exceed one (top and middle panels in **e** and **f**): p = 2.2 × 10^-20^, 2.5 × 10^-24^, 6.7 × 10^-42^, 3.0 × 10^-57^. One-side tailed t-test for whether AO significantly increased ΔF/F_0_ (bottom panels in **e** and **f**): p = 7.0 × 10^-23^, 1.5 × 10^-26^. Error bars: ± s.d. Fluorescence lifetime deconvolution along the FACED/X dimension was applied for display purposes only (**Fig. S4**); data analysis was performed on raw data.

We found that aberration correction increased the brightness of the varicosities (e.g., arrowheads, **Fig. 5c**, zoomed-in views of boxed regions in **Fig. 5b**) and glutamate transient magnitudes (**Fig. 5d**). We quantified the basal brightness (F_0_) and the glutamate transient magnitude (ΔF/F_0_) for 176 ROIs with visually evoked responses within this FOV (**Methods**). Even though aberration correction increased F_0_ by a moderate 1.2 ± 0.2 fold (mean ± s.d.; **Fig. 5e**, top panel), its impact on ΔF/F_0_ was much more drastic: For ROIs whose mean ΔF/F_0_ values during the 0-1 s period following stimulation exceeded 0.02 and thus detectable before AO correction (117 ROIs, including ROIs 59, 123, 133 in **Fig. 5d**), aberration correction increased their transient magnitudes by 2.4 ± 1.2 fold (**Fig. 5e**, middle panel); For ROIs with mean ΔF/F_0_ less than 0.02 and thus not detectable without AO (59 ROIs, including ROIs 116, 218 in **Fig. 5d**; ΔF/F_0_ of 0.001 ± 0.015), AO enhanced their ΔF/F_0_ values to 0.088 ± 0.042 (**Fig. 5e**, bottom panel).

We observed similar enhancement across four FOVs. F_0_ of 572 ROIs with visually evoked responses was increased by 1.2 ± 0.3 folds (**Fig. 5f**, top panel). For the 500 ROIs with detectable transients before aberration correction, their ΔF/F_0_ was increased by 1.7 ± 0.9 by AO (**Fig. 5f**, middle panel). For 72 ROIs with mean ΔF/F_0_ values below 0.02 in the No AO condition (ΔF/F_0_ of 0.002 ± 0.014), AO increased their ΔF/F_0_ to 0.084 ± 0.041 (**Fig. 5f**, bottom panel).

As demonstrated here, aberration correction for cranial window alone led to substantial gains in synaptic glutamate imaging *in vivo*, improving the brightness of neuronal varicosities and boosting the detectability of their visually evoked glutamate transients. Transients that were buried in noise under uncorrected conditions became clearly detectable with improved spatial resolution after AO correction.

## Conclusions

Combining AO with FACED 2PFM enabled us to achieve *in vivo* brain imaging at synapse-resolving spatial resolution and millisecond time resolution. The benefits of AO-FACED were demonstrated across diverse *in vivo* applications in the living mouse brain. Over imaging depths up to 485 μm, AO consistently enhanced fluorescence from dendritic and synaptic structures and reduced their axial profiles. In cerebral blood flow imaging, AO improved plasma signal and contrast between plasma and blood cells over large FOVs. For glutamate imaging, AO significantly increased stimulation-evoked glutamate transient magnitudes, including those that were otherwise too weak to detect, across large FOVs and hundreds of ROIs simultaneously. Together, these results establish AO-FACED 2PFM as a powerful approach of combining ultrafast *in vivo* imaging with enhanced spatial resolution at depth.

## Methods

### AO FACED 2PFM system design

We incorporated an AO module into our previously described FACED system^5–7^ (**Fig. 1a and Fig. S1**). Pulsed 1035-nm output from an ultrafast fiber laser (Monaco, 1035-40-40, Coherent) was expanded using a 2× beam expander (BE02M-B, Thorlabs) and its power was controlled by a half-wave plate (AHWP10M-980, Thorlabs) and a polarizing beam splitter (PBS253, Thorlabs). The beam was then further expanded by a 3× beam expander (BE03M-B, Thorlabs) and focused by a cylindrical lens (LJ1267RM-B, Thorlabs) on the first FACED mirror (entrance *O* on FM1 in **Fig. 1a**) of the FACED module.

The FACED module^13^ consisted of a polarizing beam splitter (CCM1-PBS253, Thorlabs), a quarter-wave plate (AQWP10M-980, Thorlabs), and a pair of FACED mirrors (250 mm in length, 30 mm in width, and 15 mm in thickness; 141031, LAYERTEC GmbH; >99.9% reflectivity in the wavelength range of 900-1050 nm, better than λ/10 flatness, <40 fs^2^ group delay dispersion per reflection). The second FACED mirror (FM2 in **Fig. 1a**) was separated by 150 mm from and slightly tilted relative to FM1 (α = 0.0125°). In this configuration, from each input pulse, 100 output pulses were generated with 1 ns temporal and 0.0125° spatial separation between adjacent pulses.

The FACED unit pupil plane^22^ (*O* or the focal plane of the cylindrical lens) was optically conjugated to a segmented deformable mirror (DM; HEX 111, Boston Micromachines Corporation) by two achromatic doublets (L1: ACT508-1000-B-ML and L2: AC508-600-AB-ML, Thorlabs). The DM surface was then conjugated to the midplane of a pair of orthogonally arranged XY galvanometers (galvos; 5 mm clear aperture, 6215H, Cambridge Technology) via another pair of achromatic doublets (L3: AC508-600-AB-ML and L4: ACT508-500-B-ML, Thorlabs), and to the back focal plane of a water-dipping objective (1.05 NA, 25×; XLPLN25XWMP2, Olympus) using a scan lens and a tube lens (L5: SL50-2P2 and L6: TTL200MP, Thorlabs). The objective was mounted on a piezo stage (SLC-1740-D-S, SmarAct) to enable axial translation. The correction collar of the objective was set to 0 for all experiments described here. In the focal plane of the objective, FACED pulses formed 100 foci spaced 0.8 µm apart and delayed by 1 ns. The fluorescence signal was collected through the same objective and directed to the detection path by a dichroic mirror (FF665-Di02-25×36, Semrock), where it passed through an emission filter (F05-525/50-25 or FF01-680/SP-25, Semrock) and was focused on a photomultiplier tube (PMT; H7422P-40, Hamamatsu) by two lenses (L7: AC508-080-A and L8: LA1951-A, Thorlabs).

During ultrafast FACED imaging, the signal from the PMT was sampled at 10 gigasamples per second using a high-speed digitizer (ADQ7DC, Teledyne SP Devices) and processed by an onboard FPGA, which averaged every 10 sampling points to represent the fluorescence signal emitted from each FACED focus (1 ns/pixel)^7^. During aberration measurement, the signal from the PMT was first sent to a low-noise current amplifier (DLPCA-200, FEMTO) and then a fast-resetting custom analog integrator, which summed the amplified photocurrent during pixel dwell time^16^. The integrated signal was then digitized by a data acquisition device (PXIe-6368, National Instruments) at 10 µs/pixel. Galvo scanning control, sample stage motion control, DM control, and synchronization among laser pulsing, scanning, and data acquisition were implemented using custom LabVIEW programs (LabVIEW 2020, National Instruments).

### Fluorescence lifetime deconvolution of AO-FACED imaging data

A machine-learning-based deconvolution method was developed to deconvolve the fluorescence signal along the FACED/X axis (**Fig. S2**). Detailed methods are described in a separate manuscript^19^. For each sample, the AO images were used to extract the fluorescence lifetime kernel, which was then applied to the aberrated images (**Fig. S4**). For biological samples used in this study, lifetime ranged from 2.5 to 3.2 ns.

### Bead samples

Carboxylate-modified fluorescent microspheres (F8826, Thermo Fisher Scientific) were immobilized on poly(l-lysine)-coated microscope slides (15-188-48, Fisher Scientific).

### Animal use

All animal experiments were conducted according to the National Institutes of Health guidelines for animal research. Procedures and protocols on mice were approved by the Institutional Animal Care and Use Committee at the University of California, Berkeley.

### Mice preparations

Virus injection and cranial window implantation procedures have been described previously^23^. Briefly, mice were anesthetized with isoflurane (Fluriso^TM^, Vet One; 1-2% by volume in oxygen) and given the analgesic buprenorphine (subcutaneously, 0.3 mg per kg of body weight). Animals were head-fixed in a stereotaxic apparatus (Model 1900, David Kopf Instruments). A 3.5-mm-diameter craniotomy was made over the left visual cortex with dura intact. A cranial window made from a coverslip (No. 1.5, Fisher Scientific) was embedded into the craniotomy and sealed in place using dental acrylic. A titanium head-post was then affixed to the skull using cyanoacrylate glue and dental acrylic. For *in vivo* structural imaging of the Thy1-GFP Line M mouse^20^, experiment was conducted on the same day of surgery under light anesthesia (0.5% isoflurane by volume in oxygen).

Some mice also underwent virus injection using a glass pipette with a 15-20 μm opening and a 45° bevel. The pipette was backfilled with mineral oil, and a fitted plunger controlled by a hydraulic manipulator (MO10, Narishige) was inserted into the pipette for loading and slowly injecting the virus to 6-12 sites (50 nl at each site) at a depth of 250 μm within the left visual cortex. For *in vivo* structural imaging of tdTomato, sparse labeling was achieved by co-injecting a 1:1 mixture of diluted AAV2/1-syn-Cre virus (original titer 3.15 × 10^13^ GC/ml, diluted 50,000-fold in phosphate buffered saline) and AAV2/1-CAG-FLEX-tdTomato (6.5 × 10^12^ GC/ml). For *in vivo* glutamate imaging, sparse expression of iGluSnFR4 was achieved using the same dilution strategy, with co-injection of diluted AAV2/1-syn-Cre virus and AAV-hSyn-flex-iGluSnFR4_v8880_NGR-WPRE (2.32 × 10^13^ GC/ml) at a 1:1 ratio. In addition, 2 µl of red beads (F8826, Thermo Fisher Scientific; 1:200,000 dilution in filtered water) were also applied to the brain surface before cranial window implantation.

*In vivo* imaging was conducted either in the awake state or under light anesthesia (0.5% isoflurane by volume in oxygen) after a recovery period of more than 3 weeks, during which animals were habituated to head fixation. Experimental details including anesthesia state, pixel size, frame rates, and post-objective power are provided in **Table S1**.

### Dye injection for blood vessel imaging

Under isoflurane anesthesia, animals were retro-orbitally injected with 50 μL of 5% (w/v) 70k molecular weight dextran-conjugated Rhodamine B fluorescent dye (D1841, Thermo Fisher Scientific).

### Visual stimulation setup for glutamate imaging

A visual stimulation program was written in MATLAB (MathWorks) using the Psychophysics Toolbox^24^ and displayed on a liquid-crystal display (P2219H, Dell Technologies). The monitor was positioned approximately 15 cm from the mouse’s right eye and oriented at ∼40° relative to the long axis of the mouse body. For glutamate imaging, each stimulation cycle consisted of a 1-s blank period, followed by a 0.05-s flash, and then a 1.95-s blank period. At each FOV, repeated trials were acquired. For the representative FOV shown in **Fig. 5**, 60 repeated trials were acquired with AO and without AO, respectively. Across four FOVs imaged (**Fig. 5f**), 44-60 trials were acquired.

### Analysis of *in vivo* structural imaging data

*In vivo* images were motion-corrected using an iterative cross-correlation method^23^. Fluorescence lifetime deconvolution along the FACED/X dimension was applied to XY MIPs (**Fig. S4**). The AO and No AO images were then compared back-to-back, with ROIs selected on the AO images and applied to the No AO images. Bicubic interpolation was applied to both XY and YZ images. The contrast of axial profiles was quantified as:

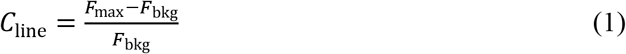

*F*_max_ represented the peak signal within the axial profile range, and *F*_bkg_ was the mean signal measured from the darkest pixels within the YZ image. The contrast ratio was then calculated as *C*_line_AO_ / *C*_line_NoAO_.

### Analysis of *in vivo* blood flow imaging data

Images were motion-corrected using an iterative cross-correlation method^23^. For each capillary segment, a line ROI was drawn along the vessel centerline, from which a kymograph was extracted from the time series. Regions of kymographs with similar stripe slopes and gaps were selected in both AO and No AO images for quantifying plasma signal and the contrast between fluorescently labeled plasma and unlabeled blood cells, by first flattening the kymographs (i.e., making the dark stripes horizontal) and then averaging along the spatial dimension to generate the line profiles in **Fig. 4g**. For each line profile, plasma signal (*F*_plasma_) was defined as the mean of the top 20% signal values of the line profile, blood cell signal (*F*_RBC_) was defined as the mean of the lowest 20% signal values of the line profile, and the plasma-cell contrast (*C*_kymo_) was defined as:

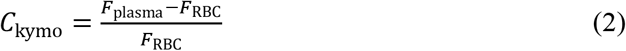

Flow velocity of the capillaries shown in **Fig. 4g** was quantified from kymographs using a cross-correlation based particle imaging velocity method^6,25^.

### Analysis of *in vivo* glutamate imaging data

Images were motion-corrected using an iterative cross-correlation method^23^. Fluorescence lifetime deconvolution was performed for display purposes only and the original data (**Fig. S4**) was used for further analysis. ROIs were manually segmented over neuronal varicosities. Pixel values within each ROI were averaged for each time frame to obtain a fluorescence trace (F). Glutamate transients were calculated as ΔF/F_0_ = (F-F_0_)/F_0_, where the basal fluorescence signal F_0_ was calculated for each trial as the average signal during the 0.5-s blank period preceding visual stimulation. Both F and ΔF/F_0_ traces were trial-averaged and temporally smoothed using a 50-ms moving window for display purposes. For glutamate transient analysis, transient magnitude ΔF/F_0_ values were defined as the mean of the trial-averaged traces within the 0-1 s window following stimulation. ROIs that had their ΔF/F_0_ values (as acquired with AO) below zero during more than 35% of time points within this window were excluded from further analysis. An ROI was classified as not visually responsive if the trial-averaged peak ΔF/F_0_ during the 0-1 s post-stimulation window was less than three times of the standard deviation (s.d.) of the baseline signal^26,27^.

## Acknowledgements

We thank Q. Zhang and D. Pan for help with AO module, W. Chen for help with LabVIEW programming, and Q. Cui for help on fluorescence lifetime deconvolution. This work was supported by the National Institutes of Health U01NS137449 and U01NS118300 (N.J.).

## Author contributions

N.J. conceived and supervised the project. J. Zhu and N.J. designed the AO-FACED module. J. Zhu built the AO-FACED module, implemented the AO control program, prepared samples, and collected and analyzed data. R.G.N. performed mouse surgeries. J. Zhong built the FACED module, wrote the data acquisition program, and assisted with data collection. I.K. designed the fluorescence lifetime deconvolution model. J. Zhu and N.J. prepared the figures and wrote the manuscript with input from all authors.

## Competing interests

N.J. and Howard Hughes Medical Institute have filed patent applications that relate to the principle of frequency-multiplexed aberration measurement. The remaining authors declare no competing interests.

## Supplementary Information

**Fig. S1.**
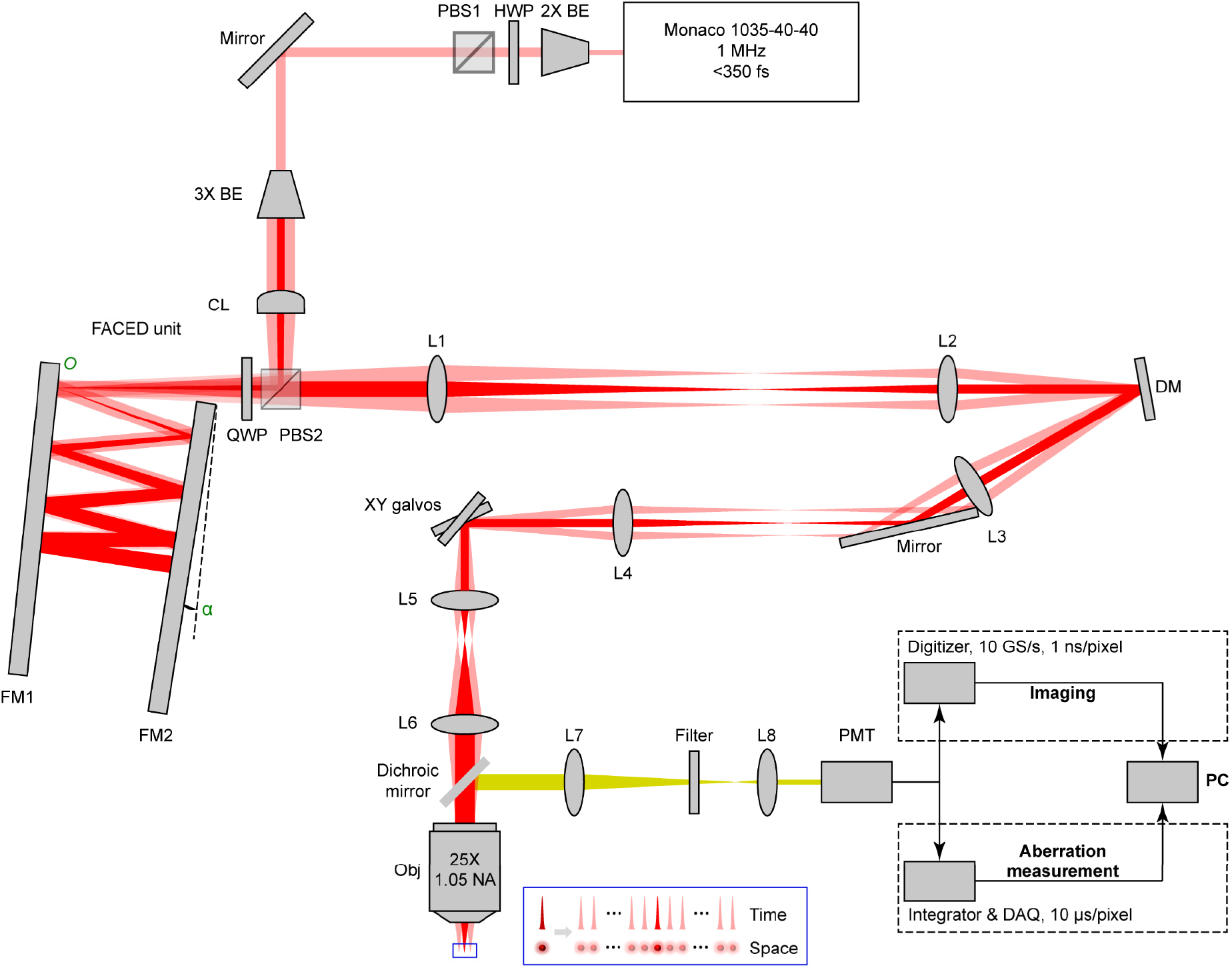
Schematic of AO-FACED 2PFM. BE: beam expander; HWP: half-wave plate; PBS: polarizing beam splitter; CL: cylindrical lens; QWP: quarter-wave plate; FM: FACED mirror; *O*: FACED beam entrance on FM1; α: relative angle between FM1 and FM2; L: lens; DM: segmented deformable mirror; Obj: objective; PMT: photomultiplier tube; DAQ: data acquisition systems; PC: personal computer. Note that the PMT signal was sampled differently during aberration measurement and FACED imaging.

**Fig. S2.**
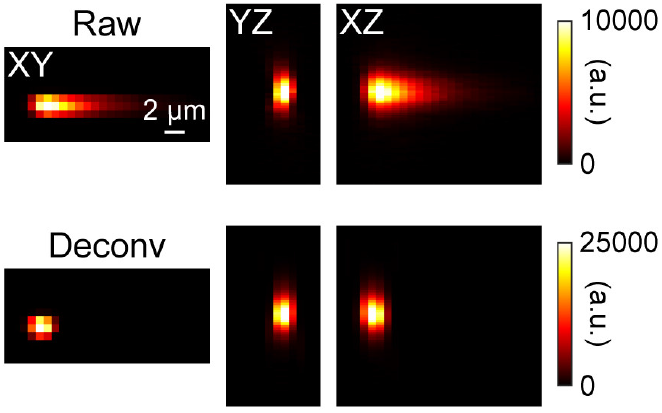
Fluorescence lifetime deconvolution of FACED images. Example lateral and axial FACED images of a 2-µm-diameter bead before (top) and after (bottom) fluorescence lifetime deconvolution along the FACED/X dimension.

**Fig. S3.**
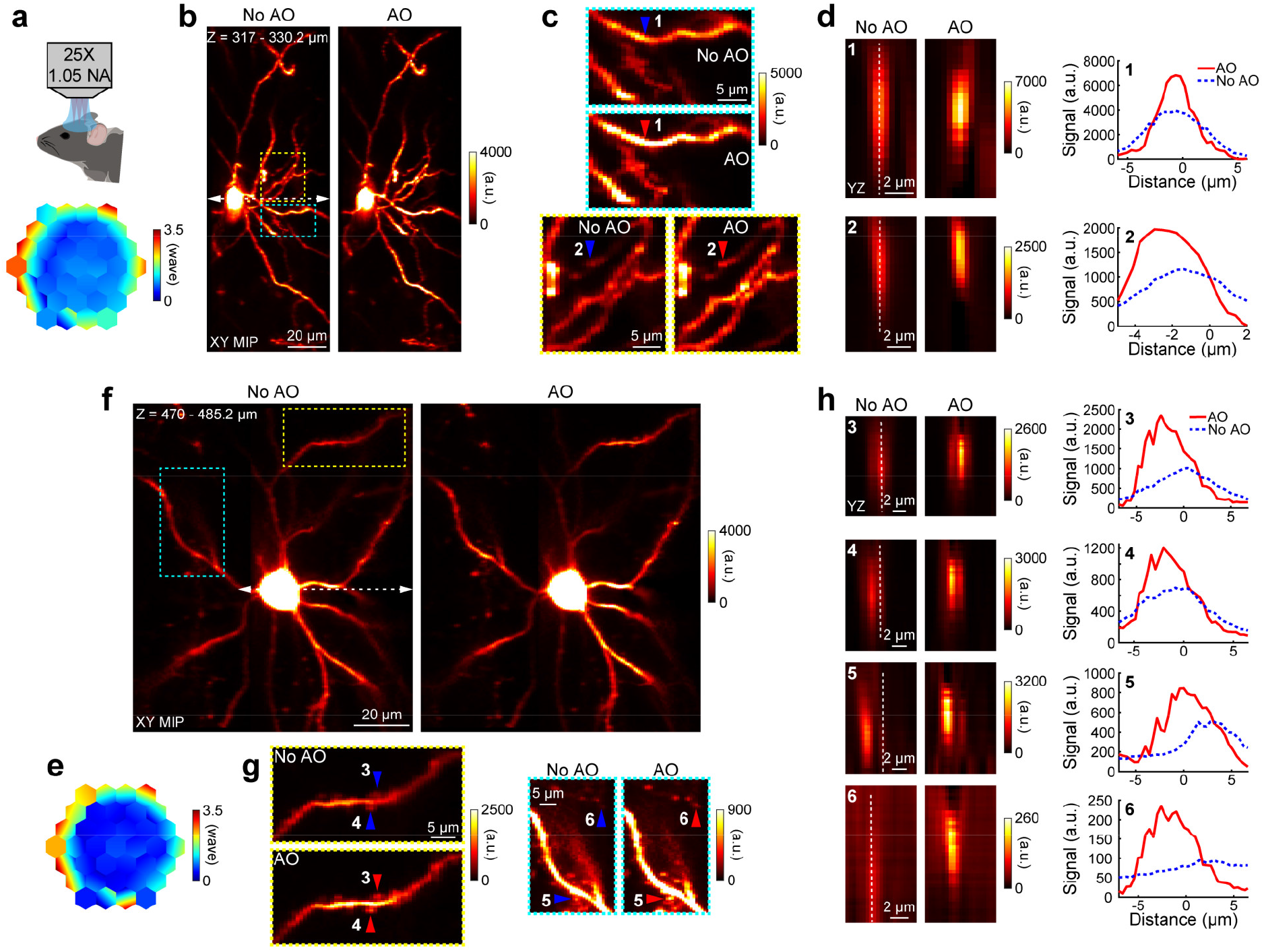
AO improves FACED 2PFM structural imaging of tdTomato-expression neurons *in vivo*. (**a**) Corrective wavefront measured from a cell body at 340 μm depth *in vivo*. (**b**) XY maximum intensity projections (MIPs) of 317-330.2 μm depth acquired without and with AO. White dashed line with arrowheads: location of FACED foci during aberration measurement. Excitation power: 76.1 mW. (**c**) Zoomed-in views of boxed regions in (**b**). (**d**) YZ images across a dendritic shaft (ROI 1) and bouton (ROI 2) (indicated by arrowheads in **c**) and their axial signal profiles along dashed white lines. (**e**) Corrective wavefront measured from a cell body at 476 μm depth *in vivo*. (**f**) XY MIPs of 470-485.2 μm depth acquired without and with AO. Excitation power: 80.2 mW. (**g**) Zoomed-in views of boxed regions in (**f**). (**h**) YZ images across dendritic shaft (ROI 3), spines (ROI 4,5), and bouton (ROI 6) (indicated by arrowheads in **g**) and their axial signal profiles along dashed white lines. Pixel size: 0.8 μm × 0.4 µm × 0.4 μm. Fluorescence lifetime deconvolution along the FACED/X dimension was applied to XY MIPs.

**Fig. S4.**
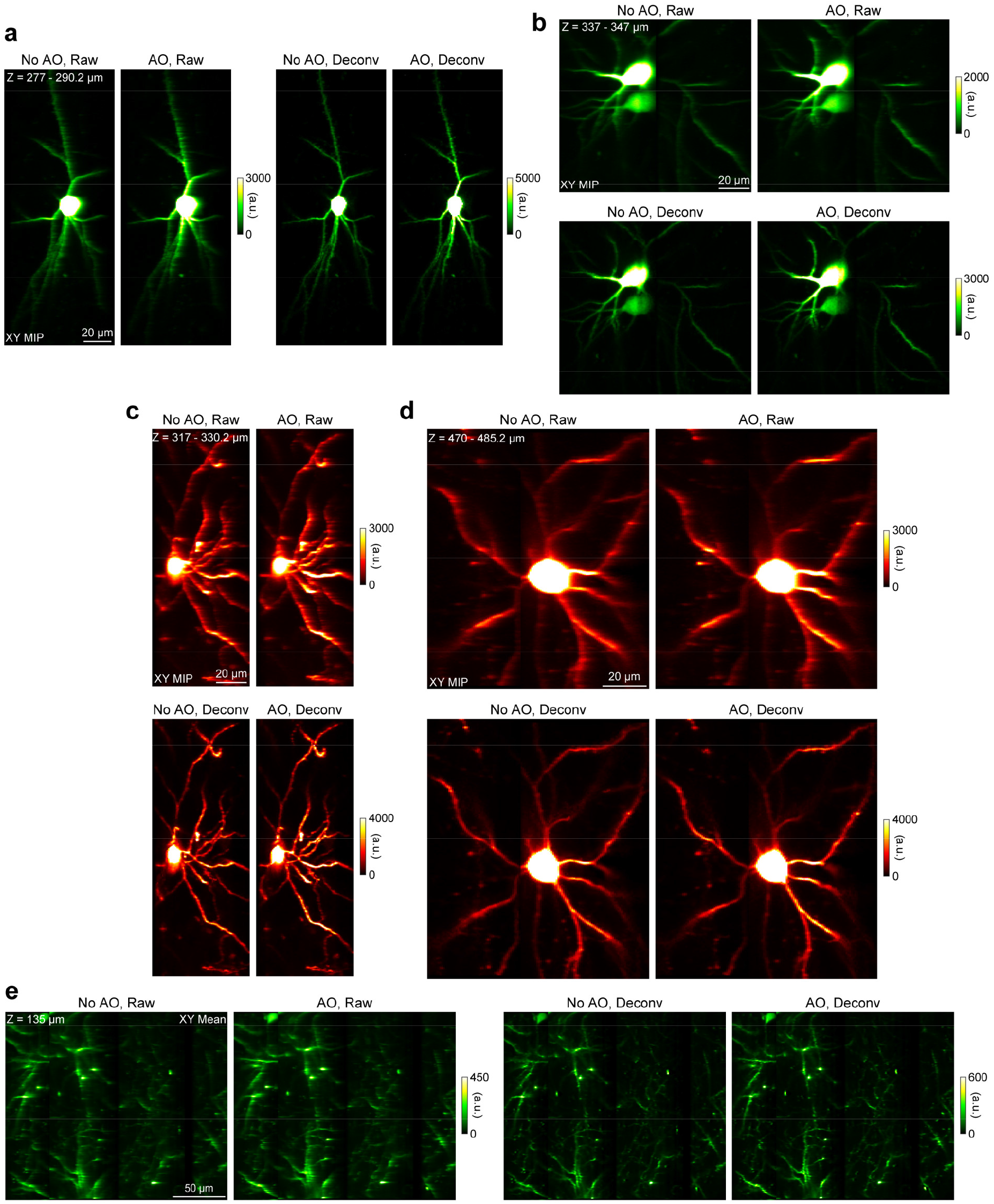
*In vivo* FACED 2PFM images before and after fluorescence lifetime deconvolution. Data correspond to *in vivo* mouse brain imaging shown in **Fig. 3b** (**a**), **Fig. 3f** (**b**), **Fig. S3b** (**c**), **Fig. S3f** (**d**), and **Fig. 5b** (**e**).

**Table S1.**
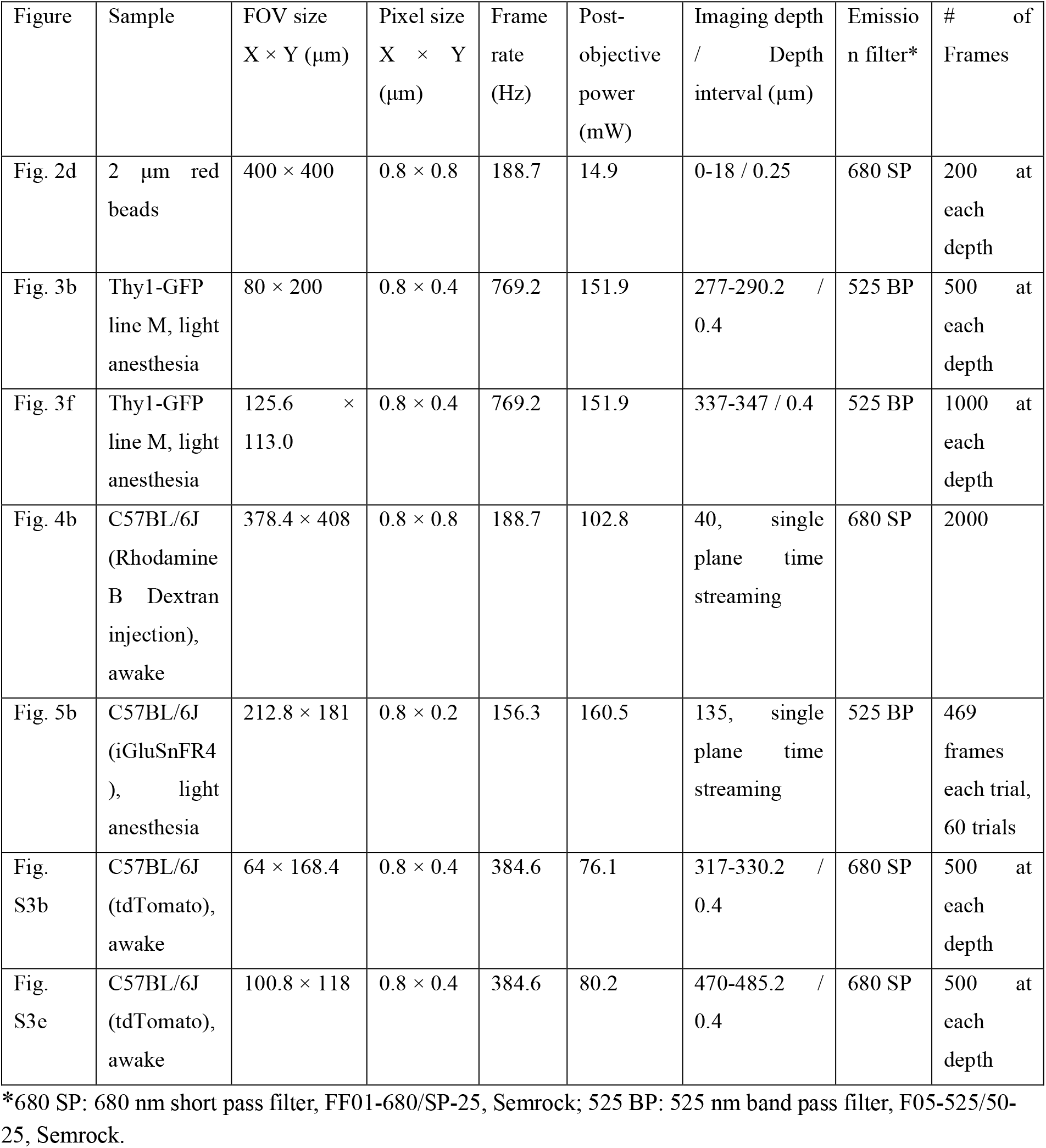
Experimental settings and image acquisition parameters.

